# Reducing Motor Variability Enhances Myoelectric Control Robustness Across Limb Positions

**DOI:** 10.1101/2023.05.05.539580

**Authors:** Simon A. Stuttaford, Matthew Dyson, Kianoush Nazarpour, Sigrid S. G. Dupan

## Abstract

**Background:** The limb position effect is a multi-faceted problem, associated with decreased upper-limb prosthesis control acuity following a change in arm position. The many factors contributing to this problem can arise from distinct environmental or physiological sources. Despite their differences in origin, the effect of each factor manifests similarly as increased variability in the detected control signal. This variability can cause incorrect decoding of user intent, leading to dropped items or inability to use the prosthesis during activities of daily living. The general approach of previous research has attempted to limit the impact of the factors or better capture the variability with data abundance. In this paper we take an alternative approach and investigate the effect of reducing the variability of the control signal by improving the consistency of muscle activity with user training.

**Methods:** Participants underwent 4 days of myoelectric training with either concurrent or delayed feedback in a single arm position. During this time, they were trained to control a two-dimensional cursor using muscles in the forearm. At the end of training participants underwent a zero feedback retention test in multiple limb positions. In doing so, we tested how well the skill learned in a single limb position generalized to untrained positions.

**Results:** We found that delayed feedback training led to more consistent muscle activity across both the trained and untrained limb positions. Analysis of patterns of activation in the delayed feedback group suggest a structured change in muscle activity occurs across arm positions. The structured changes allowed us to quantify the limb position effect by comparing trained to untrained arm positions. Different limb positions changed mean ECR and FCR muscle activity in the range of -4.3% to +18.7%. All participants were able to counter the limb position effect if given concurrent feedback, confirming our results align with existing findings.

**Conclusions:** Our results demonstrate that myoelectric user-training can lead to the retention of motor skills that are more robust to limb position changes. This work highlights the importance of reducing motor variability with practice, prior to examining the underlying structure of muscle changes associated with limb position. These findings will be useful for the majority of myoelectric prosthesis control systems and will create better quality input data leading to more robust machine-learning based prosthesis control systems.

## 1 Introduction

Multi-articulating active hand prostheses are most commonly controlled with muscle activity recorded by electromyography (EMG) sensors placed on the surface of the residual limb [1]. The information over a window of EMG data can be extracted and mapped to a prosthesis output [2]. Depending on the control scheme, this can be done with either biomimetic or physiologically distinct muscle activity [3]. Surface EMG has numerous advantages, chiefly it is non-invasive, and requires similar muscle effort to natural movement [4], which has made it ubiquitous across control schemes. However, this means that all control schemes also share the weaknesses of the approach. One challenge of using EMG is that it is a non-stationary stochastic signal -its statistical properties change over time [2]. Multiple environmental and physiological sources contribute to this nonstationarity e.g electrode conductivity or muscle fatigue; for a summary, see [5]. Transient EMG property changes can frequently occur during everyday activities, leading to unpredictable prosthesis control. Unintended activations of the prosthesis can lead to dropped items or require users to contract repeatedly until they achieve the correct grasp, fatiguing them. Limb position has been identified as one factor that can impact EMG variability [6].

The limb position effect is a term used to describe a reduction of prosthesis control acuity following a change in arm position [6]. This phenomenon is comprised of several physiological and environmental factors which combine to increase the detected signal variability [5]. Environmental factors can be defined as those that arise from contextual changes acting on the system. Whereas physiological factors include those brought about by biological or bio-mechanical reasons. Environmental factors are generally easier to diagnose and solve with better hardware design, however, physiological factors are often more challenging as they can be uncorrectable by nature. Some examples of physiological changes that affect signal variability are; muscle excitability, subcutaneous muscle displacement, and motor variability.

Fluctuations of muscle excitability can cause changes in the perceived sense of effort and force production for a given level of EMG [7, 8]. This is important because it can cause increased EMG variability for muscle contractions that are perceived to be consistent, leading to unpredictable prosthesis response. In essence, the variability induced by muscle excitability depends on several instantaneous factors, making it difficult to predict [9]. Such factors include muscle length during passive movement [10], preemptive or tonic contractions [11, 12], and the static positions of upstream proximal joints [13–15], all of which are common during activities of daily living. Furthermore, the spatial relationship between muscle and sensor is not always static. Bio-mechanical changes can cause subcutaneous muscle displacements relative to the surface electrode [16]. Certain postures or contractions may elicit changes in muscle geometry, including length, diameter as well as the relative orientation of muscle fibres [17]. These factors can, at best, alter the detected EMG features for a given muscle [18] and, at worst, record activity from a different muscle entirely [19, 20]. Thus, highly variable signals can arrive at a sensor for identical contractions. Finally, since the execution of human movement is highly over-actuated, several redundant degrees of freedom may be involved for a single coordinated movement [21]. This inherent motor abundance means there are multiple solutions for the same task goal [9,21]. Therefore, seemingly identical movements may have slightly different representations in the muscle domain. Observed differences in muscle activation between repetitions of the same movement can be partly attributed to this sensorimotor equivalence. While the variability of motor control has both positive and negative connotations with regard to motor performance [22], it is likely to have primarily negative consequences for the robustness of prosthesis control in the short-term.

Unlike the previously mentioned physiological factors, it has been demonstrated that motor variability can be lessened with practice. The reduction of which is often associated with skilled performance [22]. Training a prosthesis user to produce more consistent muscle activity is an attractive option as it is a low-cost solution for addressing input variability. However, as prostheses do not yet provide real-time feedback on the state of the user’s control signals, it is important for prosthesis users to produce skilled behavior in the absence of feedback. This is commonly referred to as retention of skill. Previous research has shown that when given real-time feedback, users are able to adapt muscle activity to counter induced perturbations in a myoelectric task space [23, 24]. However, long-term motor-learning literature suggests that this skill would not be retained as these users would be dependent on the rapid updates from real-time feedback [25–35]. Previous research has not demonstrated retention of myoelectric ability prior to testing the limb position effect. Therefore, previous results may be less certain that observed changes were due to limb position or due to input variability.

Our previous work demonstrated that appropriate myoelectric training led to the retention of skilled performance when external feedback was withdrawn [35]. This work intended to investigate if the retention of more consistent motor outputs following practice would lead to subsequent gains in robustness against the limb position effect. We trained two groups of users; one group retained their myoelectric skill, observed throughout training, during zero feedback retention tests. While the other group was highly accurate with instantaneous feedback, they did not retain this skill in the absence of feedback. After 4 days of training in a single arm position, we tested how well the retained myoelectric ability, acquired in a single position, generalized to untrained limb positions. Significant differences were found in the limb position effect between the high-retention and low-retention groups. Our research indicates that, if within-class variability can be lowered for a single arm position, the variability across other arm positions is also diminished. This suggests future research should exploit motor learning-based training prior to attempting to address the limb position effect.

## 2 Methods

### 2.1 Participants

Ten limb-intact participants (2 female, 8 male) with no known neurological or motor disorders took part in the study. All participants provided written informed consent before participating in the study, and ethical approval was granted by the local committee at Newcastle University (ref: 20-DYS-050). All participants had previously taken part in a 4 day myoelectric control experiment [35]. Data presented in this paper was acquired at the end of day 4 and on day 22.

### 2.2 Experimental setup

Participants stood 2 m in front of a 55-inch screen (Philips Q-Line DBL5530QL), with their elbow flexed at a 90° angle and their wrist in a neutral position. Eight EMG sensors (Trigno Quattro, Delsys, USA) were placed around the right forearm. Two electrodes were placed on the extensor carpi radialis (ECR) and flexor carpi radialis (FCR), located through palpation. The remaining six electrodes were placed equidistant between these electrodes. The inertial measurement unit (IMU) sensor was placed distally on the forearm. IMU data was acquired at 74 Hz, while EMG data was acquired at 2000 Hz and band-pass filtered between 20 and 450 Hz.

The muscle signals were smoothed using the mean absolute value (MAV) with a window length of 750 ms. The muscle estimation was updated at 100 Hz. As this update rate exceeds the refresh rate of the display, the participant received the visual biofeedback in the shortest possible time frame. EMG from the sensors placed on the ECR and FCR muscles were used to control the task. To normalize EMG estimates, the accompanying channels were calibrated for each participant. The calibration was performed on MAV filtered EMG data, *y*, and took place on day 1 of the experiment presented in [35]. The procedure consisted of the collection of baseline EMG activity, *y*_*r*_, and activity representing a comfortable contraction, *y*_*c*_. All subsequent tasks utilized the normalized muscle activity level, *ŷ*, for each channel:

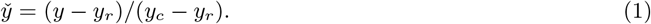

Calibration of IMU data was performed prior to each session and was repeated within the session if signal drift was detected. During the calibration, participants were asked to position their hand to nine different positions while holding their elbow in the same position. A 3×3 grid of ∼45° rotations around the elbow representing the positions is presented in Figure 1(a). After participants moved into position, quaternion data was collected and stored as a reference for the experiment. During the experiment, participants were shown where to move by highlighting the accompanying place on the grid. A filled circle on the grid denoted the current arm position. The displayed position was set to the nearest orientation stored during calibration. Positions shown in Figure 1(b) were included in the testing protocol.

**Figure 1:**
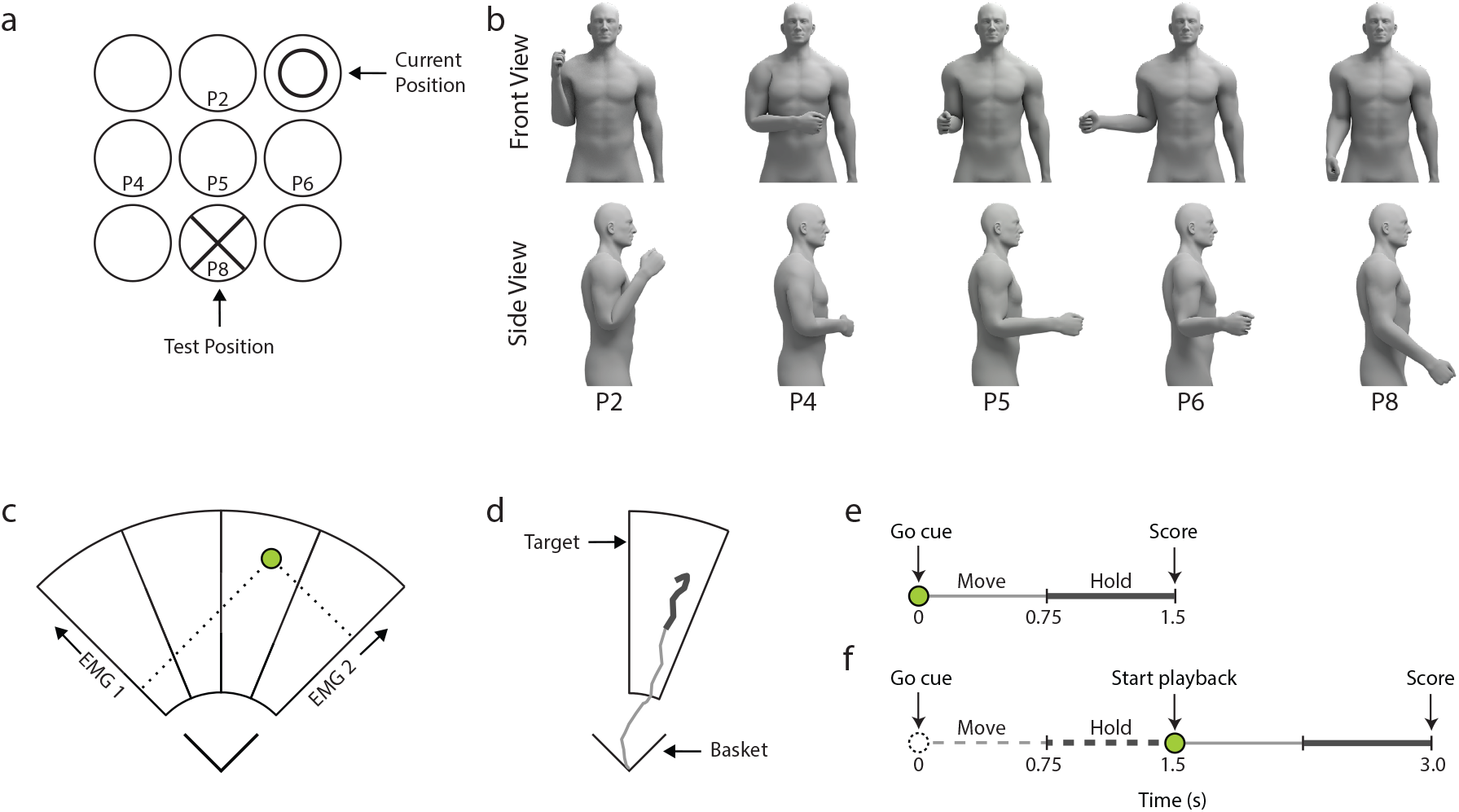
The experimental task. (a) Presented arm position widget based on inertial measurement unit (IMU) data. (b) Arm postures corresponding to the IMU widget. Feedback of the forearm direction is relative to the screen, as reflected by the IMU widget. (c) The myoelectric task interface (MCI). (d) A representative cursor trajectory during a trial. Thick cursor mark denotes the hold period. (e) Task timing structure denoting cues and the move, and hold periods for the concurrent condition. (f) Task timing structure denoting cues and the move, hold and playback periods for the delayed condition. Dashed traces correspond to the ‘blind’ control input window. Solid traces refer to the playback of the cursor’s recorded path during the move and hold periods.

### 2.3 Experimental protocol

During the experiment, participants were prompted to move their arm to a specific position, after which they performed the myoelectric control task, previously described in [35]. Briefly, participants controlled the position of a cursor within a 2-D interface which was divided into four target areas. The position of the cursor was based on the normalized muscle activity level of the ECR and FCR. The lower boundary of the interface is defined by *ŷ* = 0.3, while the upper limit is reached with *ŷ* = 1. The targets were presented when participants were at rest. Trials included consecutive move and hold periods that each lasted 750 ms. For a given trial, the score was determined by the amount of time the cursor was within or in contact with the target during the hold period.

Three feedback conditions were included in the experiment:

- Concurrent feedback: During the trial, the cursor position reflects the normalized muscle activity in real-time. The score was presented after the trial was completed (Figure 1(e)).
- Delayed feedback: The cursor was invisible during the trial. At the end of the hold period, the trajectory of the trial cursor was presented to the participant at the rate it occurred. The score was presented after the trial was completed (Figure 1(f)).
- Zero feedback: No feedback of the cursor position was provided, and no score was presented after the trial.

#### 2.3.1 Initial performance

Testing the limb position effect requires a baseline measurement, allowing for the interpretation of the performance over different positions. Therefore, participants learned the control task in position P5, with their elbow flexed at a 90° angle and their wrist in a neutral position. Results of this motor learning experiment are presented in [35].

As people are able to adapt their muscle activity on the millisecond scale based on biofeedback [36], we tested initial performance during a ‘zero feedback block’. The block was subdivided into sections, where each section was related to a specific target and consisted of 6 trials. In the first trial, participants moved into position P5, after which they completed a single trial with concurrent or delayed feedback, depending on the group condition. Then they repeated this target in positions 2, 4, 5, 6, and 8 with zero feedback. The positions were randomized within each run. Each block consisted of 12 runs, 3 for each target, and the presentation of a target for a run was randomized. At the end of day 4 participants completed 2 zero feedback blocks to test their initial performance in different limb positions. In total, each participant completed 120 zero feedback trials as part of the retention test.

#### 2.3.2 Concurrent feedback in different limb positions

A follow up session took place after an 18 day hiatus to assess long term permanency of the learned motor skill [35]. At the end of the follow-up session all participants experienced concurrent feedback and trained in multiple limb positions. This was done for two main reasons. Firstly, to investigate the effect of long-term exposure to delayed feedback, intended to prevent within-trial adaptation, on the utilization of adaptive processes within in the same task. Secondly, to assess how well participants could adapt after extensive training in a single position. Participants completed 4 blocks of 80 trials. Again, targets were presented in a serialized order. The order of presentation for each run was chosen randomly. Limb positions were randomized within a run.

#### 2.3.3 Statistical analyses

Data were visually inspected for normality and confirmed via Shapiro-Wilk tests. At the individual block level, data did not present as normal. Therefore, normally distributed data was not assumed and non-parametric statistical tests were used. Unless otherwise stated, all values reported represent the mean and standard deviation. All EMG data were inspected visually for signal artefacts. Trials which contained significant artefacts were excluded from analyses. The mean artefact rejection rate after manual inspection was 0.22% *±* 0.45%.

To test the influence of arm positions on muscle amplitude, the mean muscle activity during the hold period of the zero feedback trials was calculated for position P5 for each target. This was then used to calculate the change of the average muscle activity in the other limb positions for each target.

## 3 Results

After completing single-position testing on day 22, all participants experienced four concurrent feed-back blocks, whilst keeping their arm positioned in different postures. Figure 2 shows the average performance from the first and last of these blocks. Participants significantly improved their score across new limb positions after four blocks of concurrent feedback training (First, *Mdn* = 0.71; Last, *Mdn* = 0.90; Wilcoxon signed-rank, *p <* 0.01). No significant differences were found for comparisons between groups on the first or last concurrent feedback blocks.

**Figure 2:**
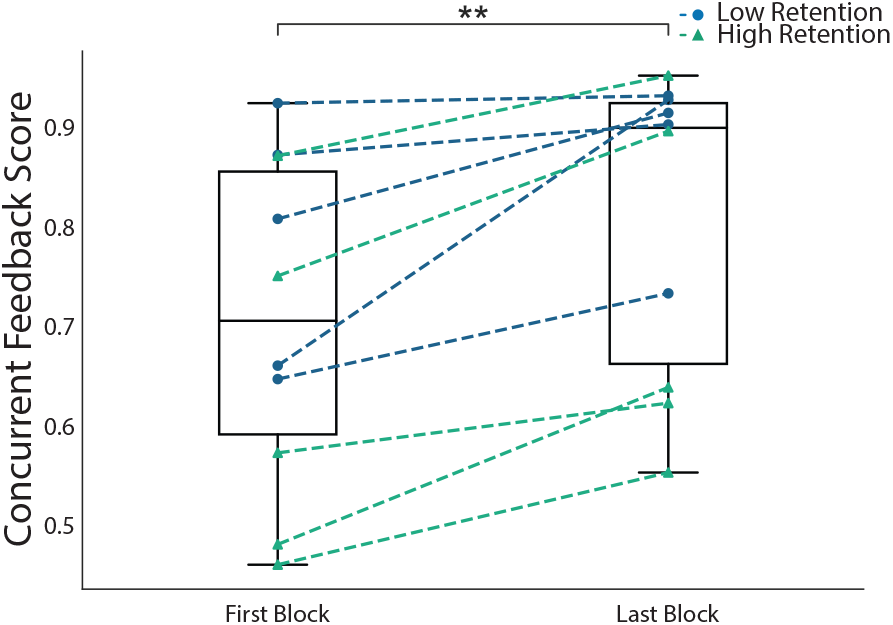
Learning to counter the limb position effect with concurrent feedback. Box plots correspond to the mean scores across all participants. Only untrained arm position trials are included. Centroid lines, medians; solid boxes, interquartile ranges; whiskers, overall ranges. Dashed lines correspond to individual participant mean scores. Asterisks denote level of statistical significance (Wilcoxon signed-rank, *p <* 0.01).

Figure 3 shows multi-position retention collected on day 4. At the end of day 4, participants completed two retention blocks consisting of trials carried out under multiple untrained arm positions. Figure 3 shows data from the multiple arm position retention blocks. There was a significant difference for position P5(low-retention, 0.28 *±* 0.12; high-retention, 0.54 *±* 0.17; *p <* 0.05). No significant differences were found for positions P2 (low-retention, 0.31 *±* 0.11; high-retention, 0.45 *±* 0.16), P4 (low-retention, 0.29 *±* 0.13; high-retention, 0.48 *±* 0.15), P6 (low-retention, 0.25 *±* 0.17; high-retention, 0.45 *±* 0.22), or P8 (low-retention, 0.26 *±* 0.12; high-retention, 0.51 *±* 0.18).

**Figure 3:**
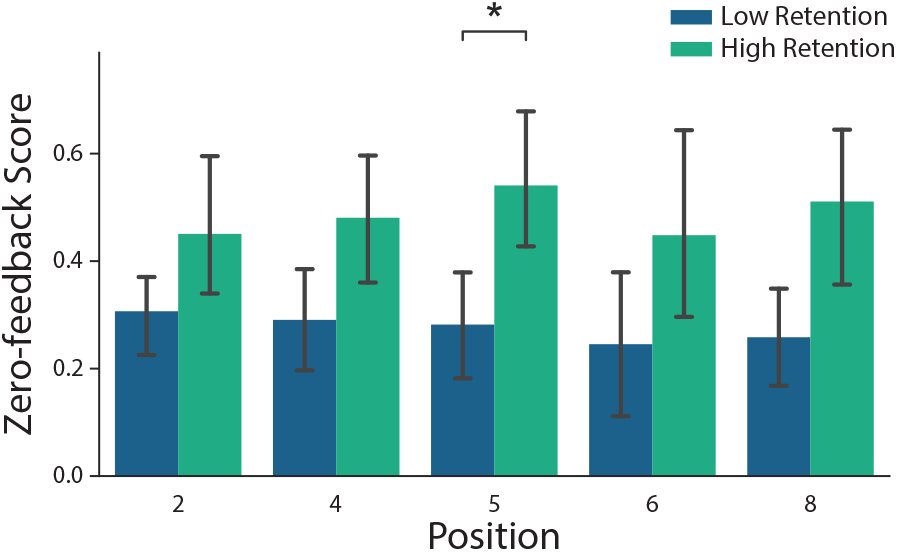
Baseline retention between groups. zero feedback scores across limb positions pre-multi-position training. Bars represent the mean. Error bars represent 95% confidence intervals. Asterisk refers to significant differences (*p <* 0.05).

Figure 4 illustrates the impact of limb position on cursor location during the retention test. The center of each ellipse is the mean cursor coordinate during the hold period over the zero feedback trials. The semi-major and semi-minor axes represent the standard error of the mean for the cursor’s location in the direction of each control muscle. In general, the high-retention group had less variable muscle activity in both control sensors than the low-retention group. In addition, the within-group variability between limb positions was lower in the high-retention group. Table 1 shows the area of the ellipses, which are proportional to the product of the standard errors of each control sensor. Comparing the effect of each limb position between groups shows the total variability in the high-retention group was consistently lower than the low-retention group for every target.

**Figure 4:**
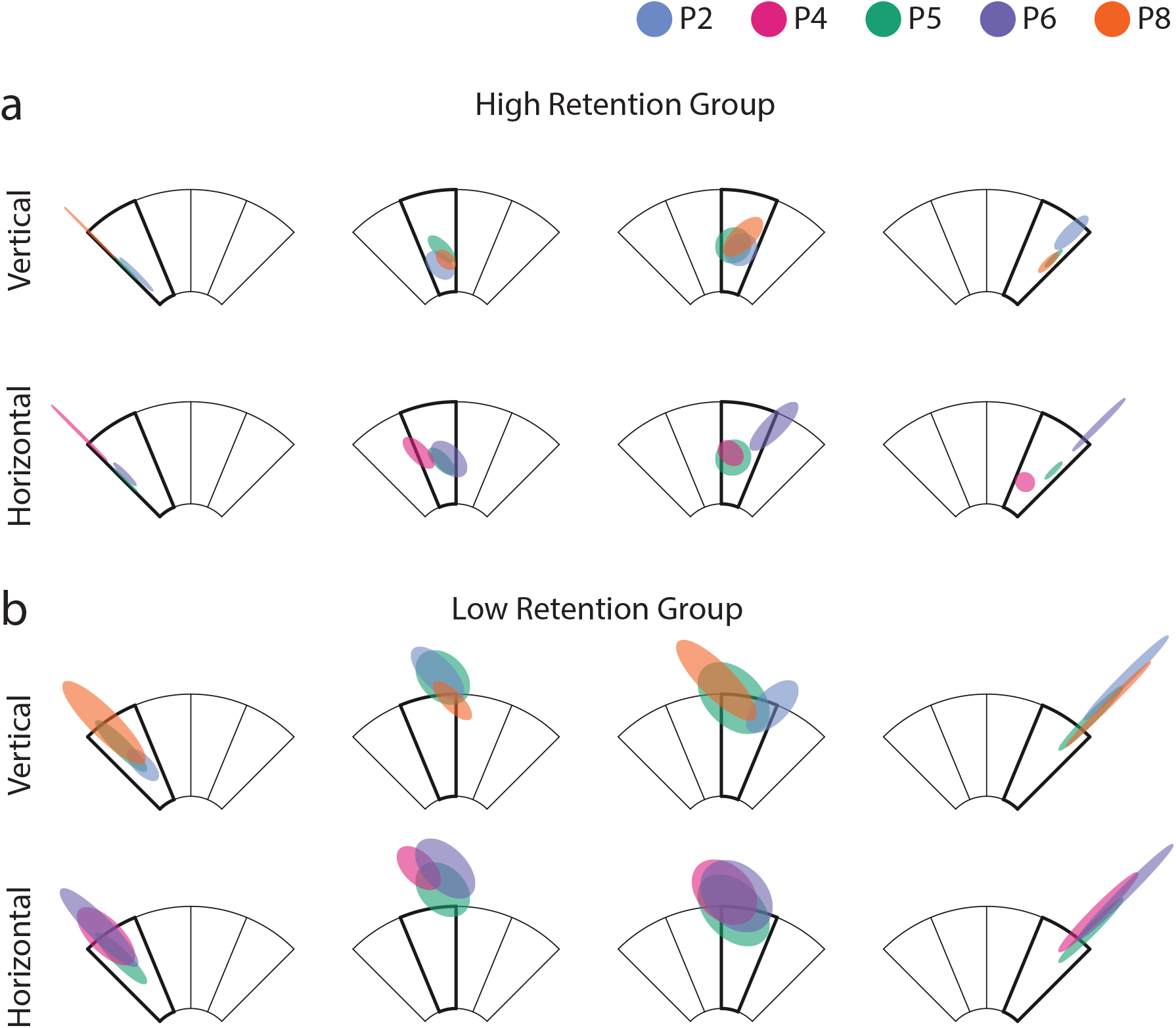
Group differences in muscle activity variability during the retention test. (a) and (b) show the cursor’s location across limb positions for the high-retention and low-retention feedback groups, respectively. Plotted activity corresponds to zero feedback trials of the retention blocks. Rows refer to arm positions located in the vertical (P2, P6) and horizontal (P6, P8) planes, relative to the torso. Position P5 is plotted across both planes for comparison. Columns separate the goal target presented during the hold period, shown in bold. Ellipse centre represents the group’s mean muscle activation during the hold period. Ellipse semi-major and semi-minor axes represent the standard error in the corresponding control muscle.

**Table 1:** Area of ellipses representing the standard error, presented in Figure 4. Values are factored by *π ·* 10^−3^*units*^2^.

|  |  | Low Retention Group |  |  |  | High Retention Group |  |  |  |
| --- | --- | --- | --- | --- | --- | --- | --- | --- | --- |
|  |  | Target: 1 | 2 | 3 | 4 | 1 | 2 | 3 | 4 |
| Position | 2 | 8.2 | 23.4 | 24.1 | 13.9 | 2.1 | 8.0 | 11.7 | 7.1 |
|  | 4 | 27.7 | 18.1 | 45.4 | 16.3 | 2.0 | 7.8 | 6.8 | 4.1 |
|  | 5 | 12.4 | 31.6 | 53.6 | 10.0 | 0.5 | 5.7 | 14.2 | 1.2 |
|  | 6 | 28.8 | 35.0 | 56.3 | 16.7 | 1.5 | 12.7 | 15.4 | 4.4 |
|  | 8 | 36.9 | 10.8 | 40.9 | 8.6 | 1.1 | 3.7 | 13.3 | 2.2 |

Figure 4 shows that some ellipses cross target boundaries, which represents a greater misclassification rate. For positions in both vertical and horizontal planes relative to the torso, there is a relatively large amount of overlap into adjacent targets for the low-retention group. Whereas, for the high-retention group, instances of misclassification mainly occur in the positions located in the horizontal plane. Because any underlying patterns within the low-retention group’s data are more likely to be obscured by their greater variability, subsequent analyses focus on findings from the high-retention group only.

Figure 5(a) investigates the relative changes of ECR and FCR muscle activity of high-retention group participants, in different arm positions compared to position P5. Activity of the ECR muscle increased when the elbow was flexed in position P2 (10.8 *±* 14.2%), and when the shoulder was rotated outward in P6 (18.7*±*9.4%), relative to P5. Whereas, comparatively less change of ECR activity was observed for positions P4 (−2.8 *±* 5.8%) and P8 (−0.9 *±* 8.6%). Conversely, mean FCR activity increased when the shoulder was rotated inwards in position P4 (13.8 *±* 1.6%), and when the elbow was extended in P8 (5.6 *±* 14.4%), relative to P5. Whilst relatively smaller changes were observed for positions P2 (−4.3 *±* 4.6%) and P6 (2.4 *±* 7.8%).

**Figure 5:**
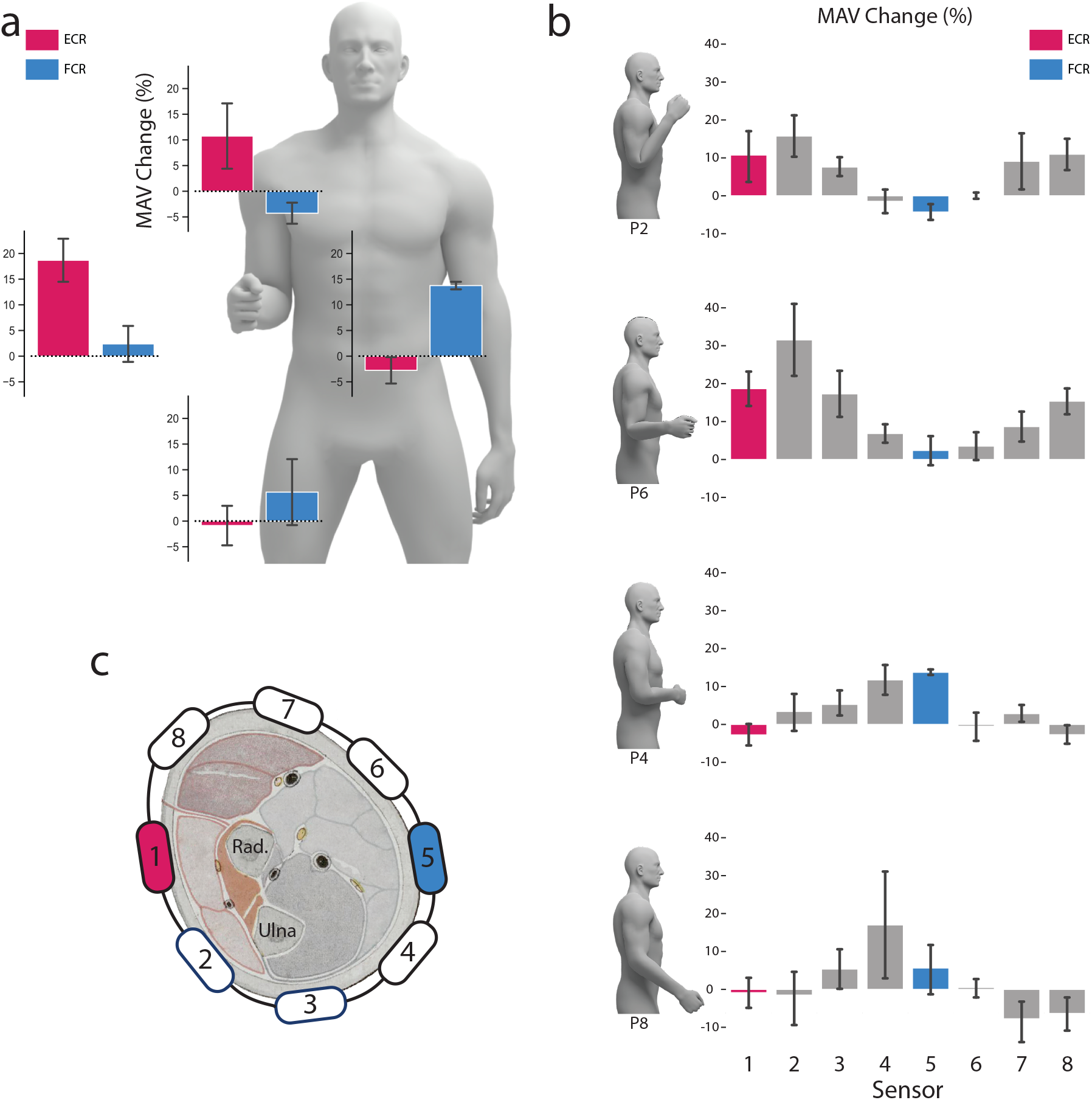
The effect of arm position on the high-retention group’s muscle activity. (a) ECR and FCR activity changes compared to position P5. The location of each bar plot around the human model reflects results from the corresponding arm position. (b) Activity changes compared to position P5 across all recording sites. (c) Stylistic representation of electrode positions around the forearm. Forearm cross section is adapted from [37] which is in the public domain. Bars represent means. Error bars represent the standard error of the mean.

Figure 5(b) shows the relative change of muscle activity across all recording sites relative to position P5. In general, a similar change in muscle activity occurs across recording sites for positions P4 and P6. For both positions, the peak of the profile occurs at sensor 1, with the trough at sensor 4. In addition, a similar pattern of muscle activity changes can also be observed for positions P4 and P8. Peak activity occurs between sensors 4 and 3, respectively. However, the variability across sensors for position P8 is comparatively high between participants.

## 4 Discussion

The drop in control performance following a change in posture has challenged the robustness of myoelectric prostheses and continues to be the focus of much research [38]. Mitigation attempts in the pattern recognition domain have focused on acquiring high-dimensional data to provide more tolerant decision boundaries [5]. However, these studies did not investigate the influence of user practice prior to addressing the limb position effect. While previous research in motor learning-based control schemes has leveraged user learning to counter limb position-induced perturbations [23, 24], there is no evidence that this can be done without real-time feedback. In this study, we compared the impact of limb position between two groups after long-term myoelectric training. The group that trained with delayed feedback retained less variable muscle activity during the zero feedback tests. The other group received real-time feedback, which resulted in higher muscle activity variability. Our findings are threefold. Firstly, delayed feedback training in a single-arm position lead to the retention of muscle contraction patterns that were more robust against limb position changes. Secondly, by training participants to produce more consistent muscle activity, we could more confidently attribute observed activation patterns to the limb position effect. Finally, our results stress the importance of providing appropriate feedback mechanisms during both training and real-world use of prosthetic devices.

To counter the limb position effect, pattern recognition solutions have attempted to capture more variability to provide better decision boundaries [5], but have not attempted to reduce the motor variability inherent in the data presented to the classifier. The applicability of our results to pattern recognition are easier to interpret by considering the targets in the myoelectric task as analogous to decision boundaries. For classification to be robust, motor activity must be sufficiently consistent that the EMG signal remains quasi-stationary, i.e. within decision boundaries. In our case, however, rather than adapting the decision boundaries to fit user data, as is done during classifier training, the boundaries are pre-defined and the user must learn to adapt their input accordingly. Following this analogy, Figure 4 illustrates the impact of appropriate myoelectric training on within-class variability. When compared to concurrent feedback, we can see that delayed feedback training in one limb position reduces within-class variability across all limb positions. Reducing within-class variability can, in turn, increase class separability [39]. This is visually demonstrated in Figure 4 by less overlap with other targets. In the context of pattern recognition, the difference between the low-retention and high-retention groups suggests user training could be beneficial for enhancing the consistency of contraction patterns and hence improve classification robustness. Previous studies have found that increased class separability achieved during pattern recognition training does not necessarily transfer to prosthesis control [40]. However, these studies did not investigate retention. We have previously shown the importance of demonstrating retention of myoelectric ability prior to demonstrating transfer [35, 41].

It has been shown that some EMG features are highly reproducible when contraction intensity is controlled [42–45]. However, this is distinct from how prosthesis users modulate muscle activity during real-world use, as their production of force typically lacks precise external feedback and is comparatively unconstrained. Removing external feedback can impact the quality of EMG and has been shown to lead to the generation of higher amplitude patterns [46]. Our results suggest that delayed feedback training is beneficial for improving the consistency of contraction intensity. Reducing contraction variability enabled greater clarity of the underlying structure of the limb position effect. As such, analysis focused on the high-retention group, which showed they had retained more consistent muscle activity than the low-retention group. Although reduced variability in one position generalized to untrained positions (Figure 4), limb positions in the horizontal plane produced results which suggests a more structured change (Figure 5). This was further investigated by analysing changes in the recorded muscle activity.

In general, we see that ECR activity increases when the arm is lifted or moved to the right, while positions down or across the body increase activity in FCR. One possible reason for the increase in EMG relative to position P5 could be due to shared muscle excitability patterns between certain postures. Equally, similar patterns of subcutaneous muscle displacements may be shared between position P2, P6 as well as P4 and P8. Furthermore, for each pair of arm positions, a similar pattern seems to hold for the activity profile across all eight sensors (Figure 5(b)). It is important to note that the changes in activity recorded across sensors may not directly correspond to changes in the underlying physiology. For example, activity from a muscle can contaminate the signal recorded by adjacent surface EMG sensors due to crosstalk [19, 47]. Increased activity from a single muscle can lead to a subsequent increase in the contribution of that signal arriving at nearby sensors. It is difficult to ascertain the contribution of muscle-electrode displacement or muscle excitability with surface EMG sensors. In reality, both factors may act simultaneously, and the contribution of each may differ across positions that share seemingly similar EMG responses. Furthermore, due to the spatial resolution limits of surface EMG, we cannot be certain of the underlying pattern of any induced changes. However, irrespective of the exact physiological structure, these results represent the signal changes that would be detected by the electrodes and presented to a prosthesis decoder.

This study intended to investigate muscle activity patterns following changes in limb position. This was only possible because we first conducted a retention study. This enabled greater certainty that any observed patterns were due to limb position and not inconsistent muscle activity. As the participants who trained with delayed feedback reduced their control signal variability, we were able to show that the reduced variability observed in one position also reduced the variability in untrained positions. Finally, we wanted to know how correctable the perturbations induced by the limb positions were, and thus all participants experienced concurrent feedback at the end of the experiment. We found that, when given concurrent feedback, both groups were able to adapt to limb position changes. In addition, the ability to utilize this information and counter perturbations induced by limb position improved over blocks. This is in agreement with existing literature [23, 24]. As the main focus of this study was to investigate if reduced muscle activity variability in one position could generalize to untrained positions, we did not set up an additional long-term, delayed feedback, retention study. Future research should look at the effectiveness of delayed feedback training in multiple limb positions. Previous research has shown that concurrent adaptation may not transfer to real-world control, as users would be dependent on the real-time feedback they were trained with [25–35]. One theory of why this occurs is that frequent feedback may overwhelm attention and disrupt learning processes that lead to retention. For example, real-time biofeedback may encourage rapid ad-hoc corrections that prevent a stable skill from forming. Whereas, less frequent or less informative feedback might encourage focus to shift towards proprioceptive signals as a source of guidance. Thus leading to retention of skill when external feedback mechanisms are removed. This is important to consider when reporting the performance of participants with real-time feedback. Until prostheses widely provide high fidelity, real-time feedback of the users’ control signals, it is essential for motor skill to be produced in the absence of feedback.

One limitation of this study is the exclusive use of limb-intact participants. Previous work has found that limb different participants’ control signals are less affected by limb position. This is thought to be due to anatomical differences. For example, a shorter residual limb enacts a smaller moment of inertia during movement or limb stabilisation [48]. Therefore less compensatory muscle activity is needed to maintain certain postures. Furthermore, anchoring of the muscle during amputation may also contribute to reduced muscle displacement during movement for some participants [49–51]. These characteristics diminish the overall variability of muscle activity between arm positions. Therefore, we expect our results with limb-intact participants to reflect a more extreme case of induced physiological perturbations. Furthermore, the findings did not take into account the secondary effects of arm dynamics on the loading and movement of a prosthesis socket. However, this study intended to investigate the impact of reducing the variability introduced by the central nervous system rather than environmental or contextual factors, which have been studied previously. Finally, it is not immediately clear how well user training is likely to scale to machine learning-based systems which require high dimensional EMG data. Future work will investigate whether user training produces comparable results in machine learning-based systems as it does in motor learning-based systems.

## Acknowledgements

This work has been supported by the Engineering and Physical Sciences Research Council (EPSRC) via grants EP/R511584/1 and EP/R004242/2, the National Institute for Health Research (NIHR) Award ID: NIHR201310, and Science Foundation Ireland under grant 21/PATH-S/9605.

## References

1. G. Hefftner and G. Jaros. The electromyogram EMG as a control signal for functional neuromuscular stimulation-part II: practical demonstration of the EMG signature discrimination system. IEEE Trans. Biomed. Eng., 35(4):238–242, 1988.

2. M. Zecca, S. Micera, M. C. Carrozza, and P. Dario. Control of multifunctional prosthetic hands by processing the electromyographic signal. Crit. Rev. Biomed. Eng., 30(4-6):459–485, 2002.

3. A. Fougner, Ø. Stavdahl, P. J. Kyberd, Y. G. Losier, and P. A. Parker. Control of upper limb prostheses: Terminology and proportional myoelectric control—a review. IEEE Trans. Neural Syst. Rehabilitation Eng., 20(5):663–677, 2012.

4. P. Parker, K. Englehart, and B. Hudgins. Myoelectric signal processing for control of powered limb prostheses. J. Electromyogr. Kinesiol., 16(6):541–548, 2006.

5. I. Kyranou, S. Vijayakumar, and M. S. Erden. Causes of performance degradation in non-invasive electromyographic pattern recognition in upper limb prostheses. Front. Neurorobot., 12:58, 2018.

6. A. Fougner, E. Scheme, A. D. C. Chan, K. Englehart, and Ø. Stavdahl. Resolving the limb position effect in myoelectric pattern recognition. IEEE Trans. Neural Syst. Rehabilitation Eng., 19(6):644–651, 2011.

7. F. Gelli, F. Del Santo, R. Mazzocchio, and A. Rossi. Force estimation processing as a function of the input–output relationship of the corticospinal pathway in humans. Eur. J. Neurosci., 22(5):1127–1134, 2005.

8. F. Del Santo, F. Gelli, F. Ginanneschi, T. Popa, and A. Rossi. Relation between isometric muscle force and surface emg in intrinsic hand muscles as function of the arm geometry. Brain Res., 1163:79–85, 2007.

9. M. T. Turvey, R. E. Shaw, and W. Mace. Issues in the theory of action: Degrees of freedom, coordinative structures and coalitions. In Attention and performance VII, pages 557–595. Routledge, 1978.

10. L. Chye, K. Nosaka, L. Murray Edwards, and G. Thickbroom. Corticomotor excitability of wrist flexor and extensor muscles during active and passive movement. Hum. Mov. Sci., 29(4):494–501, 2010.

11. B. Andersen, K. M. Rosler, and M. Lauritzen. Nonspecific facilitation of responses to transcranial magnetic stimulation. Muscle & Nerve: Official Journal of the American Association of Electrodiagnostic Medicine, 22(7):857–863, 1999.

12. P. M. Rossini, D. Burke, R. Chen, L. G. Cohen, Z. Daskalakis, R. Di Iorio, V. Di Lazzaro, F. Ferreri, P. B. Fitzgerald, M. S. George, et al. Non-invasive electrical and magnetic stimulation of the brain, spinal cord, roots and peripheral nerves: Basic principles and procedures for routine clinical and research application. an updated report from an IFCN committee. Clin. Neurophysiol., 126(6):1071–1107, 2015.

13. F. Dominici, T. Popa, F. Ginanneschi, R. Mazzocchio, and A. Rossi. Cortico-motoneuronal output to intrinsic hand muscles is differentially influenced by static changes in shoulder positions. Exp. Brain Res., 164(4):500–504, 2005.

14. F. Ginanneschi, F. Dominici, A. Biasella, F. Gelli, and A. Rossi. Changes in corticomotor excitability of forearm muscles in relation to static shoulder positions. Brain Res., 1073:332–338, 2006.

15. J. P. M. Mogk, L. M. Rogers, W.M. Murray, E. J. Perreault, and J. W. Stinear. Corticomotor excitability of arm muscles modulates according to static position and orientation of the upper limb. Clin. Neurophysiol., 125(10):2046–2054, 2014.

16. A. Rainoldi, M. Nazzaro, R. Merletti, D. Farina, I. Caruso, and S. Gaudenti. Geometrical factors in surface emg of the vastus medialis and lateralis muscles. J. Electromyogr. Kinesiol., 10(5):327–336, 2000.

17. L. Mesin, M. Joubert, T. Hanekom, R. Merletti, and D. Farina. A finite element model for describing the effect of muscle shortening on surface emg. IIEEE. Trans. Biomed. Eng., 53(4):593–600, 2006.

18. L. Mesin, S. Smith, S. Hugo, S. Viljoen, and T. Hanekom. Effect of spatial filtering on crosstalk reduction in surface emg recordings. Med. Eng. Phys., 31(3):374–383, 2009.

19. J. P. M. Mogk and P. J. Keir. Crosstalk in surface electromyography of the proximal forearm during gripping tasks. J. Electromyogr. Kinesiol., 13(1):63–71, 2003.

20. M. Yung and R. P. Wells. Changes in muscle geometry during forearm pronation and supination and their relationships to EMG cross-correlation measures. J. Electromyogr. Kinesiol., 23(3):664–672, 2013.

21. N. A. Bernstein. The co-ordination and regulation of movements. Oxford: Pergamon Press, 1967.

22. K. M. Newell and D. M. Corcos. Variability and motor control, volume 491, pages 1–12. Human Kinetics Publishers Champaign, IL, 1993.

23. J. M. Hahne, S. Dähne, H.J. Hwang, K. R. Müller, and L.C. Parra. Concurrent adaptation of human and machine improves simultaneous and proportional myoelectric control. IEEE Trans. Neural Syst. Rehabilitation Eng., 23(4):618–627, 2015.

24. J. M. Hahne, M. Markovic, and D. Farina. User adaptation in myoelectric man-machine interfaces. Sci. Rep., 7:4437, 2017.

25. A. W. Salmoni, R. A Schmidt, and C. B. Walter. Knowledge of results and motor learning: a review and critical reappraisal. Psychol. Bull., 95(3):355–386, 1984.

26. D. E. Sherwood. Effect of bandwidth knowledge of results on movement consistency. Percept. Mot. Ski., 66(2):535–542, 1988.

7. S. P. Swinnen, R. A. Schmidt, D. E. Nicholson, and D. C. Shapiro. Information feedback for skill acquisition: Instantaneous knowledge of results degrades learning. J. Exp. Psychol. Learn. Mem. Cogn., 16(4):706–716, 1990.

8. C. J. Winstein and R. A. Schmidt. Reduced frequency of knowledge of results enhances motor skill learning. J. Exp. Psychol. Learn. Mem. Cogn., 16(4):677–691, 1990.

9. R. A Schmidt. Frequent augmented feedback can degrade learning: Evidence and interpretations. Tutorials in Motor Neuroscience, pages 59–75, 1991.

30. D. E. Young and R. A. Schmidt. Augmented kinematic feedback for motor learning. J. Mot. Behav., 24(3):261–273, 1992.

31. C. J. Winstein, P. S. Pohl, and R. Lewthwaite. Effects of physical guidance and knowledge of results on motor learning: Support for the guidance hypothesis. Res. Q. Exerc. Sport., 65(4):316–323, 1994.

32. R. A Schmidt and G. Wulf. Continuous concurrent feedback degrades skill learning: Implications for training and simulation. Hum. Factors, 39(4):509–525, 1997.

33. J. H. Park, C. H. Shea, and D. L. Wright. Reduced-frequency concurrent and terminal feedback: A test of the guidance hypothesis. J. Mot. Behav., 32(3):287–296, 2000.

34. D. Maslovat, K. M. Brunke, R. Chua, and I. M. Franks. Feedback effects on learning a novel bimanual coordination pattern: Support for the guidance hypothesis. J. Mot. Behav., 41(1):45–54, 2009.

35. S. A. Stuttaford, S. S. G. Dupan, K. Nazarpour, and M. Dyson. Delaying feedback during pre-device training facilitates the retention of novel myoelectric skills: a laboratory and home-based study. J. Neural Eng., 2023. https://doi.org/10.1088/1741-2552/acc4ea doi:10.1088/1741-2552/acc4ea.

36. M. Couraud, D. Cattaert, F. Paclet, P. Y. Oudeyer, and A. de Rugy. Model and experiments to optimize co-adaptation in a simplified myoelectric control system. J. Neural Eng., 15(2):026006, 2018.

37. H. Braus, C. Elze, and R.B. Green. Anatomie des Menschen: ein Lehrbuch für Studierende und Ärzte. Julius Springer, Berlin, 1921.

38. E. Campbell, A. Phinyomark, and E. Scheme. Current trends and confounding factors in myoelectric control: Limb position and contraction intensity. Sensors, 20(6):1613, 2020.

39. B. Hudgins, P. Parker, and R.N. Scott. A new strategy for multifunction myoelectric control. IEEE Trans. Biomed. Eng., 40(1):82–94, 1993.

40. M.B. Kristoffersen, A.W. Franzke, R.M. Bongers, M. Wand, A. Murgia, and C.K van der Sluis. User training for machine learning controlled upper limb prostheses: a serious game approach. J. NeuroEng. Rehabil., 18(1):1–15, 2021.

41. S. S. G. Dupan, S. A. Stuttaford, K. Nazarpour, and M. Dyson. Transfer of abstract control skills to prosthesis use. In Proc. Myoelectric Controls Symp. (MEC), pages 94–97, 2022.

42. E. R. Buskirk and P. V. Komi. Reproducibility of electromyographic measurements with inserted wire electrodes and surface electrodes. Acta Physiologica Scandinavica, 79(2):29A, 1970.

43. T. Moritani and H. A. Devries. Reexamination of the relationship between the surface integrated electromyogram (iemg) and force of isometric contraction. Am. J. Phys. Med., 57(6):263–277, 1978.

44. T. Moritani and H. A. Devries. Neural factors versus hypertrophy in the time course of muscle strength gain. Am. J. Phys. Med., 58(3):115–130, 1979.

45. T. Moritani and M. Muro. Motor unit activity and surface electromyogram power spectrum during increasing force of contraction. Eur. J. Appl. Physiol. Occup. Physiol., 56(3):260–265, 1987.

46. M. B. Kristoffersen, A. W. Franzke, C. K. van der Sluis, A. Murgia, and R. M. Bongers. The effect of feedback during training sessions on learning pattern-recognition-based prosthesis control. IEEE Trans. Neural Syst. Rehabilitation Eng., 27(10):2087–2096, 2019.

47. C. J. De Luca and R. Merletti. Surface myoelectric signal cross-talk among muscles of the leg. Electroencephalogr. Clin. Neurophysiol., 69(6):568–575, 1988.

48. S. H. Roy, S. L. Wolf, and D. A. Scalzitti. The Rehabilitation Specialist’s Handbook, page 1013. F. A. Davis Co., Philadelphia, PA, 4 edition, 2012.

49. Y. Geng, P. Zhou, and G. Li. Toward attenuating the impact of arm positions on electromyography pattern-recognition based motion classification in transradial amputees. J. Neuroeng. Rehabilitation., 9:1–11, 2012.

50. Y. Teh and L. J. Hargrove. Understanding limb position and external load effects on real-time pattern recognition control in amputees. IEEE Trans. Neural Syst. Rehabilitation Eng., 28(7):1605–1613, 2020.

51. N. Jiang, S. Muceli, B. Graimann, and D. Farina. Effect of arm position on the prediction of kinematics from emg in amputees. Med. Biol. Eng. Comput., 51:143–151, 2013.

